# Homeoprotein neuroprotection of embryonic neuronal cells

**DOI:** 10.1101/695684

**Authors:** Stephanie E. Vargas Abonce, Mélanie Leboeuf, Alain Prochiantz, Kenneth L. Moya

## Abstract

Most homeoprotein transcription factors have a highly conserved internalization domain used in intercellular transfer. Internalization of homeoproteins ENGRAILED1 or ENGRAILED2 promotes the survival of adult dopaminergic cells, whereas that of OTX2 protects adult retinal ganglion cells. Here we characterize the *in vitro* neuroprotective activity of several homeoproteins in response to H_2_O_2_. Protection is observed with ENGRAILED1, ENGRAILED2, OTX2, GBX2 and LHX9 on midbrain and striatal embryonic neurons whereas cell-permeable c-MYC shows no protective effects. Therefore, five homeoproteins belonging to 3 different classes (ANTENNAPEDIA, PAIRED and LIM) share the ability to protect embryonic neurons from midbrain and striatum. Because midbrain and striatal neurons do not express the same repertoire of the 4 proteins, a lack of neuronal specificity together with a general protective activity can be proposed. In contrast, hEN1 and GBX2 exerted no protection on non-neuronal cells including mouse embryo fibroblasts, macrophages or HeLa cells. For the 4 proteins, protection against cell-death correlated with a reduction in the number of H_2_O_2_-induced DNA break foci in midbrain and striatal neurons. In conclusion, within the limit of the number of cell types and homeoproteins tested, homeoprotein protection against oxidative stress-induced DNA breaks and death is specific to neurons but shows no homeoprotein or neuronal type specificity.

**SIGNIFICANCE STATEMENT:** Homeoproteins are DNA binding proteins regulating gene expression throughout life. Many of them transfer between cells and are thus internalized by live cells. This has allowed for their use as therapeutic proteins in animal models of Parkinson disease and glaucoma. Part of their therapeutic activity is through a protection against neuronal death. Here we show that internalized homeoproteins from three different classes protect embryonic ventral midbrain and striatal neurons from oxidative stress, both at the level of DNA damage and survival. The interest of this finding is that it lends weight to the possibility that many homeoproteins play a role in neuroprotection through shared mechanisms involving, in particular, DNA protection against stress-induced breaks.

## INTRODUCTION

Homeoprotein (HP) transcription factors, discovered on the basis of their developmental functions, remain expressed in the adult where they exert not fully understood physiological activities (Di Nardo et al., 2018). Several HPs transfer between cells thanks to highly conserved secretion and internalization domains present in their DNA-binding site or homeodomain (HD). HP internalization has allowed for the use of OTX2, ENGRAILED1 (EN1) and ENGRAILED2 (EN2) (collectively ENGRAILED), as therapeutic proteins in animal models of Parkinson disease (ENGRAILED) and glaucoma (OTX2) (Sonnier et al., 2007; Alvarez-Fischer et al., 2011; Torero-Ibad et al., 2011; Thomasson et al., 2019).

The two proteins EN1 and EN2 are expressed in adult midbrain dopaminergic (mDA) neurons (Di Nardo et al., 2007). These neurons degenerate progressively in Parkinsonian patients, in classical Parkinson disease animal models and in the *EN1*-heterozygous mouse. In all models tested, ENGRAILED injected or infused is internalized by mDA neurons and prevents their death, even following a strong and acute oxidative stress provoked by a 6-hydroxydopamine hydrobromide (6-OHDA) injection at the level of the Substantia Nigra pars compacta (SNpc) (Rekaik et al., 2015). The mechanisms involved in this protection have started to be analyzed. ENGRAILED internalization stimulates translation of complex I mitochondrial proteins, restores the chromatin epigenetic marks disrupted by the stress and allows for DNA repair as quantified by the number of γH2AX foci (Alvarez-Fischer et al., 2011; Rekaik et al., 2015). In addition, ENGRAILED represses the expression of LINE-1 mobile elements caused by oxidative stress *in vitro* and *in vivo* (Blaudin de Thé et al., 2018). Because of the epigenetic nature of the protection mechanisms, a single injection of ENGRAILED has long-lasting effects, including in non-human primates (Thomasson et al., 2019), opening the way for a therapeutic use of this HP.

In view of developing ENGRAILED as a therapeutic protein, human EN1 (hEN1) was produced and purified and an assay was adapted to test hEN1 for neuroprotection against oxidative stress and, in particular, to evaluate protein activity, specificity and stability. Because OTX2 has a similar survival effect on mDA neurons and retinal ganglion cells (Rekaik et al, 2015; Torero-Ibad et al., 2011), it could be that protection against oxidative stress is a shared property of several HPs with little HP and/or neuronal specificity. To test this hypothesis, the protective effect of EN1, EN2, OTX2, GBX2 and LHX9 was evaluated on midbrain and striatal neurons in culture **as well as testing hEN1 on a neuronal cell line**. We show that the 5 proteins, but not cell-permeable c-MYC, protect embryonic midbrain and striatal neurons against oxidative stress-induced cell death and DNA damage and cell death induced by hydrogen peroxide (H_2_O_2_) but are ineffective on mouse embryo fibroblasts (MEFs), peritoneal macrophages and HeLa cells. **hEN1 was also protective against 6-OHDA in the dopaminergic Lhumes cells**. The protective activity **in the different cell types and against the two stressors** involves reducing DNA damage.

## MATERIAL AND METHODS

### Animal treatment

All animals were treated in accordance with the guide for the care and use of laboratory animals (US National Institutes of Health), the European Directive number 86/609 (EEC Council for Animal Protection in Experimental Research and Other Scientific Utilization) and with French authorizations n° 00703.01 and APAFIS #6034-2016071110167703 v2.

### Cell cultures

For neuronal primary cultures, pregnant Swiss mice (Janvier) were sacrificed by cervical dislocation 14.5 days post-conception (dpc) and the embryos were extracted and placed in Phosphate Buffer Saline /Glucose 0.6% (PBS-glucose). Striatal or midbrain structures were dissected in 2 mL of PBS-glucose and cells were mechanically dissociated and plated in previously coated in Poly-L-ornithine (1/100) and 2.5μg/ml laminin (Sigma) 96 wells plates for LDH assay and 24 well plates with glass coverslips for immunocytochemistry. Cells were cultured in Neurobasal medium (Life Technologies) supplemented with glutamine (500 µM, Sigma), glutamic acid (3.3 mg/l Sigma) aspartic acid (3.7 mg/l, Sigma), anti-anti (Gibco) and B27 (Gibco) (NB+) for 24 hours at 37°C in a humidified incubator with 5% CO_2_ atmosphere. All experiments were performed at 6 days in vitro (DIV).

Primary mouse embryo fibroblasts (MEFs) were isolated from the skin of 11 dpc Swiss mouse embryos (Janvier) according to Jozefczuk, et al. (2012). Cells were grown on 75 cm^2^ tissue culture flask at 37°C in a humidified incubator with 5% CO_2_ atmosphere. Cells at 80% confluence were detached using 0.05% Trypsin-EDTA (Gibco), plated at a density of 12500 cells per well in 96 well tissue cuture plastic plates and cultured for 24h in Dulbecco’s Modified Eagle Medium (DMEM), high glucose, GlutaMAX (Gibco) supplemented with 10% (v/v) fetal bovine serum (FBS) (Gibco) before the addition of 10 μM cytosine arabinoside (Ara-C) from Sigma.

HeLa cells were maintained in DMEM, 1g/L D-Glucose L-Glutamine, Pyruvate (Gibco) supplemented with 10% (v/v) FBS (Gibco). Cells were grown on 75 cm^2^ tissue culture flask at 37°C in a humidified incubator with 5% CO_2_ atmosphere. Cells at 80% confluence were detached using 0.05% Trypsin-EDTA (Gibco) and plated at a density of 12500 cells in each well in 96 well tissue cuture plastic plates. Cells were cultured for 24h before stopping proliferation with 10 μM Ara-C.

Macrophages were isolated from the mouse peritoneal cavity of eight week-old Swiss female mice (Janvier). Mice were sacrificed by cervical dislocation and peritoneal washes were performed using Hank’s solution (HBSS). After massaging the peritoneum, the fluid containing resident macrophages was collected, seeded and plated at a density of 100000 cells per well in 96 well tissue cuture plastic plates in DMEM + GlutaMAX (Gibco) 2% FBS (Gibco) at 37°C in a humidified incubator with 5% CO^2^ atmosphere.

### LUHMES cell culture

LUHMES (ATCC® CRL-2927) cells were thawed rapidly at 37°C, transferred to a 15mL Falcon tube with 3 mL of AdvDMEM and centrifuged for 7 minutes at 190*g*. Supernatant was discarded and 1mL of DMEM was added to the pellet. After gentle resuspension, the cells were placed in AdvDMEM+FGF (40ng/mL) and cultured for 3 days at 37°C before trypsinization (0.025%Trypsine-0.1g/L EDTA in PBS) for 5 minutes at 37°C, followed by the addition of 4mL of AdvDMEM medium and centrifugation for 7 minutes at 190*g*. The cells were dissociated with 1mL of AdvDMEM+FGF and plated on previously coated in laminin (1μg/mL) and Poly-L-Ornithine (50μg/mL) 96 wells plates for the LDH assay, or coated in laminin (1μg/mL) and Poly-L-Ornithine (500μg/mL) glass coverslips for immunocytochemistry. Cells were cultured in Advanced DMEM+FGF for 1 day or 3 days at 37°C in a humidified incubator with 5% CO_2_ atmosphere for the LDH assay or γ-H2AX foci analysis, respectively. Proteins ware added at the times indicated in the text in the presence 1μM Ara-C.

### RT-qPCR

Total RNA was extracted using the RNeasy Mini kit (Qiagen) and reverse transcribed using the QuantiTect Reverse Transcription kit (Qiagen). The RT-qPCR was made using SYBR-Green (Roche Applied Science) and Light Cycler 480 (Roche Applied Science). Data were analyzed using the « 2-ddCt » method and values were normalized to Glyceraldehyde 3-phosphate dehydrogenase (*Gapdh*).

### Protein production

Chicken ENGRAILED2 (chEN2) and mutant chicken ENGRAILED2 (SR-EN2), mouse EN1 (mEN1), human EN1 (hEN1) and mouse OTX2 (mOTX2) were prepared as described (Joliot et al., 1998; Torero et al., 2011). Cell-permeable recombinant human c-MYC was purchased from Abcam (ab169901) and human GBX2 (hGBX2) and LHX9 (hLHX9) were purchased from Proteogenix. Endotoxins were removed by phase separation according to Aida and Pabst, 1990. Unless stated otherwise, proteins were stored at -20°C.

### Protein treatment and oxidative stress

Cell were incubated with different concentrations of HPs diluted in culture media. For neutralization, HPs were preincubated with a 10-fold molar excess of antibody for 1 hour at 37°C. For LDH and trypan blue assay, oxidative stress was induced by incubation for 2 hours at 37°C in 50mM H_2_O_2_ (Sigma Aldrich) **or 6-OHDA (6-hydroxydopamine hydrobromide; 300**, **200 or 100** μ**M)** diluted in culture media. For DNA break analysis, H_2_O_2_ (100μM) **or 6-OHDA (10 or 50** μ**M)** were added for 1 hour. For dye-exclusion survival analysis, the media was replaced with 0.16% trypan blue for 5 min at room temperature then replaced with PBS and the number of cells excluding or not trypan blue were counted blind in five fields of view at 20x, five wells per condition. The LDH assay was carried out using the CytoTox 96® Non-Radioactive Cytotoxicity Assay (Promega) according to manufacturer instructions.

### Immunocytochemistry

Coverslips were washed three times in PBS, fixed in 4% paraformaldehyde for 30 minutes at room temperature (RT), washed in PBS 3 times, permeabilized with PBS/0.5% Triton (Sigma) for 45 min at RT and placed in 100mM Glycine for 30 min at RT. After a 1 hour incubation at RT in PBS/10% Natural Goat Serum (NGS, Invitrogen)/1% Triton, primary antibodies were added overnight at 4°C. The next day coverslips were washed 3 times in PBS, incubated with secondary antibodies for 2 hours at RT, washed 3 times in PBS and mounted in DAPI Fluoromount-G® (Southern Biotech). The mouse monoclonal anti-γH2AX antibody (IgG1) is from clone JBW301 (Millipore) and the mouse monoclonal anti-ß-Tubulin III (IgG2A) is from clone SDL.3D10 (Sigma). Both antibodies were used at a 1/500 dilution. Alexa-conjugated goat anti-mouse antibodies (Life Technologies) were used at a 1/2000 dilution.

### Quantification of DNA damage

Images corresponding to a coverslip diameter were acquired with a Nikon i90 microscope and exported to ImageJ. DNA damage was quantified by counting the number of γH2AX foci present in the nucleus of Tuj1 positive cells. Counting was carried out blind to conditions on four coverslips per condition.

### Statistical analysis

Data are expressed as mean ± SD if not otherwise indicated and results analyzed with Prism v6 GraphPad). For the trypan blue experiment, statistical significance was determined by one-way ANOVA and two-tailed t-test using five wells per condition. For the LDH assay experiments statistical significance was determined by one-way ANOVA and a *post hoc* Dunnett’s test for comparisons to H_2_O_2_ using 8 replicates per condition. (*p < 0.5, **p < 0.005, ***p ≤ 0.001 and ****p < 0.001 in all experiments). Statistical power for each significant difference was determined using the statistical power calculator (https://www.stat.ubc.ca/~rollin/stats/ssize/n2.html on 18-19 February, 2019 and on 4 July, 2109).

**Table 1.** Statistical power analysis

| Data structure |  | Type of test | Power |
| --- | --- | --- | --- |
| Trypan Blue | Control vs H<sub>2</sub>O<sub>2</sub> | Post-hoc t-test,<br>two-tailed | 1.00 |
|  | H<sub>2</sub>O<sub>2</sub> vs 12.5 nM |  | 0.92 |
| Figure 1-A | H<sub>2</sub>O<sub>2</sub> vs 12.5 nM | Dunnett's multiple<br>comparisons | 1.000 |
|  | H<sub>2</sub>O<sub>2</sub> vs 2.5 nM |  | 1.000 |
|  | H<sub>2</sub>O<sub>2</sub> vs 25 pM |  | 1.000 |
|  | H<sub>2</sub>O<sub>2</sub> vs 12.5 pM |  | 1.00 |
|  | H<sub>2</sub>O<sub>2</sub> vs 25 pM |  | 1.000 |
|  | H<sub>2</sub>O<sub>2</sub> vs 1.25 nM |  | 1.000 |
| Figure 1-B | H<sub>2</sub>O<sub>2</sub> vs 12.5 nM | Dunnett's multiple<br>comparisons | 1.000 |
|  | H<sub>2</sub>O<sub>2</sub> vs 2.5 nM |  | 1.00 |
| Figure 1-C | H<sub>2</sub>O<sub>2</sub> vs 12.5 nM 1x frozen | Dunnett's multiple<br>comparisons | 1.000 |
|  | H<sub>2</sub>O<sub>2</sub> vs 2.5 nM 1x frozen |  | 1.00 |
|  | H<sub>2</sub>O<sub>2</sub> vs 12.5 nM 5x frozen |  | 1.00 |
|  | H<sub>2</sub>O<sub>2</sub> vs 2.5 nM 5x frozen |  | 1.000 |
|  | H<sub>2</sub>O<sub>2</sub> vs 12.5 nM 4°C |  | 1.000 |
|  | H<sub>2</sub>O<sub>2</sub> vs 2.5 nM 4°C |  | 1.000 |
| Figure 1-D | H<sub>2</sub>O<sub>2</sub> vs 12.5 nM chEn2 | Dunnett's multiple<br>comparisons | 1.000 |
|  | H<sub>2</sub>O<sub>2</sub> vs 2.5 nM chEn2 |  | 1.000 |
|  | H<sub>2</sub>O<sub>2</sub> vs 12.5 nM hEn1Q50A |  | 1.000 |
|  | H<sub>2</sub>O<sub>2</sub> vs 2.5 nM hEn1Q50A |  | 1.000 |
| Figure 2-A | H<sub>2</sub>O<sub>2</sub> vs 12.5 nM hEn1 | Dunnett's multiple<br>comparisons | 1.000 |
|  | H<sub>2</sub>O<sub>2</sub> vs 2.5 nM hEn1 |  | 1.000 |
|  | H<sub>2</sub>O<sub>2</sub> vs 12.5 nM mOtx2 |  | 1.000 |
|  | H<sub>2</sub>O<sub>2</sub> vs 2.5 nM mOtx2 |  | 1.000 |
|  | H<sub>2</sub>O<sub>2</sub> vs 12.5 nM hGbx2 |  | 1.000 |
|  | H<sub>2</sub>O<sub>2</sub> vs 2.5 nM hGbx2 |  | 1.000 |
|  | H<sub>2</sub>O<sub>2</sub> vs 12.5 nM hLhx9 |  | 1.000 |
| Figure 2-B | H<sub>2</sub>O<sub>2</sub> vs 12.5 nM hEn1 | Dunnett's multiple<br>comparisons | 1.000 |
|  | H<sub>2</sub>O<sub>2</sub> vs 2.5 nM hEn1 |  | 1.000 |
|  | H<sub>2</sub>O<sub>2</sub> vs 12.5 nM mOtx2 |  | 1.000 |
|  | H<sub>2</sub>O<sub>2</sub> vs 2.5 nM mOtx2 |  | 1.000 |
|  | H<sub>2</sub>O<sub>2</sub> vs 12.5 nM hGbx2 |  | 1.000 |
|  | H<sub>2</sub>O<sub>2</sub> vs 2.5 nM hGbx2 |  | 1.000 |
|  | H<sub>2</sub>O<sub>2</sub> vs 12.5 nM hLhx9 |  | 1.000 |
|  | H<sub>2</sub>O<sub>2</sub> vs 2.5 nM hLhx9 |  | 1.000 |
| Figure 3-B | H<sub>2</sub>O<sub>2</sub> vs 2.5 nM mEn1 | Dunnett's multiple comparisons | 1.00 |
|  | H<sub>2</sub>O<sub>2</sub> vs 1.25 nM mEn1 |  | 1.000 |
|  | H<sub>2</sub>O<sub>2</sub> vs 0.62 nM mEn1 |  | 1.00 |
|  | H<sub>2</sub>O<sub>2</sub> vs 0.31 nM mEn1 |  | 1.00 |
|  | H<sub>2</sub>O<sub>2</sub> vs 3.3 nM mOtx2 |  | 1.000 |
|  | H<sub>2</sub>O<sub>2</sub> vs 1.65 nM mOtx2 |  | 1.00 |
|  | H<sub>2</sub>O<sub>2</sub> vs 0.82 nM mOtx2 |  | 1.000 |
|  | H<sub>2</sub>O<sub>2</sub> vs 0.41 nM mOtx2 |  | 1.00 |
|  | H<sub>2</sub>O<sub>2</sub> vs 2.68 nM hGbx2 |  | 1.000 |
|  | H<sub>2</sub>O<sub>2</sub> vs 1.34 nM hGbx2 |  | 1.000 |
|  | H<sub>2</sub>O<sub>2</sub> vs 0.67 nM hGbx2 |  | 1.000 |
|  | H<sub>2</sub>O<sub>2</sub> vs 0.33 nM hGbx2 |  | 1.000 |
|  | H<sub>2</sub>O<sub>2</sub> vs 2.27 nM hLhx9 |  | 1.000 |
|  | H<sub>2</sub>O<sub>2</sub> vs 1.14 nM hLhx9 |  | 1.000 |
|  | H<sub>2</sub>O<sub>2</sub> vs 0.56 nM hLhx9 |  | 1.000 |
|  | H<sub>2</sub>O<sub>2</sub> vs 0.28 nM hLhx9 |  | 1.000 |
| Figure 3-C | H<sub>2</sub>O<sub>2</sub> vs Stavudine | Dunnett's multiple comparisons | 1.000 |
|  | H<sub>2</sub>O<sub>2</sub> vs hEn1 |  | 1.000 |
| Figure 3-D | H<sub>2</sub>O<sub>2</sub> vs Stavudine | Dunnett's multiple comparisons | 1.000 |
|  | H<sub>2</sub>O<sub>2</sub> vs hEn1 |  | 1.000 |
| Figure 4-A | -hEN1 vs +hEN1 at 300uM 6-OHDA | Post-hoc t-test, two-tailed | 1.000 |
|  | -hEN1 vs +hEN1 at 200uM 6-OHDA |  | 1.000 |
| Figure 4-B | -hEN1 vs +hEN1 at 10uM 6-OHDA | Post-hoc t-test, two-tailed | 1.000 |
|  | -hEN1 vs +hEN1 at 50uM 6-OHDA |  | 1.000 |

## RESULTS

### EN1 protects embryonic neurons against oxidative stress

Five independent hEN1, one mEN1 (mouse) and one chEN2 (chicken) preparations produced similar results. Protective activity was measured by the LDH cytotoxicity assay, except for one experiment in which trypan blue exclusion was used for comparison. In the trypan blue experiment, mean survival of E14.5 ventral midbrain cells in the control condition was 89.3 ± 4.0%. Two hours after adding H_2_O_2,_ neuron survival was reduced to 36.5 + 14.1% (p<0.0001 compared with Control). The survival of neurons pretreated with 12.5nM hEN1 was significantly greater (66.4 ± 9.6%, p<0.005) compared to H_2_O_2_-treated cells. The LDH assay gave qualitatively similar results and was thus used thereafter, making it easier to test different preparations and dose-responses over a large range of HP concentrations.

LDH is a cytosolic enzyme and this assay measures LDH released into the culture medium after the lysis of live cells. Figure 1A shows a hEN1 dose-response survival experiment for embryonic midbrain neurons at 6 DIV. Two hours after oxidative stress with 50mM H_2_O_2_, more than 90% of the cells are dead. Pretreatment of the cells with 1.25pM to 12.5nM significantly increases their survival from about 28 to 86% in an EN1 dose-dependent manner. It is of note that H_2_O_2_ effects were variable between experiments with oxidative stress-induced cell death varying between 50 and 90%. Based on this dose-response, 12.5 and 2.5nM HP concentrations allowing for total or near total protection were used in further experiments.

**Figure 1.**
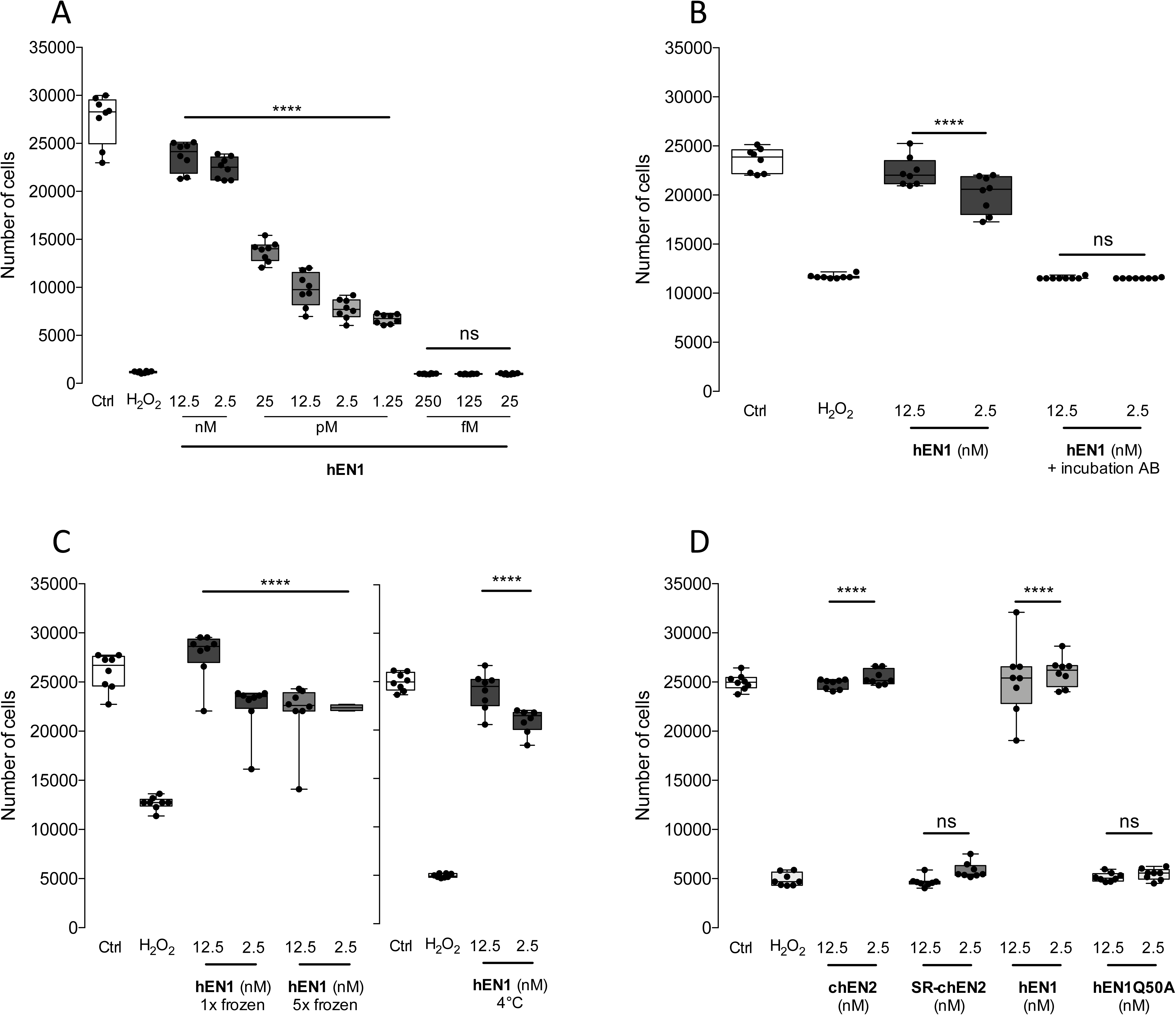
ENGRAILED protection of embryonic midbrain neurons. A: hEN1 dose-dependent survival of embryonic neurons after H_2_O_2_ oxidative stress. B: Preadsorption of hEN1 with and anti-ENGRAILED antibody abrogates hEN1 neuroprotection. C: hEN1 submitted to repeated freeze-thaw cycles (left) or maintained at 4°C for six weeks (right) has significant neuroprotective activity against oxidative stress. D: ENGRAILED internalization and high affinity DNA binding are necessary for ENGRAILED neuroprotection.

Since hEN1 is a recombinant protein purified from bacterial extracts, its activity could be in part due to a contaminant. As shown in Figure 1B, protective activity was fully abolished by pre-incubation (1 hour at 37°C) of the protein with an anti-EN1 polyclonal antibody (Alvarez-Fischer et al., 2011) establishing that the neuroprotective activity is entirely due to hEN1. To examine hEN1 stability, aliquots were frozen on dry ice and thawed once or five times. Midbrain neurons were treated with hEn1 and 24h later stressed with 50mM H_2_O_2_. Human EN1 frozen and thawed once provided 100% protection and 84 to 70% if frozen and thawed five times (Fig1C, left). The protein maintained at 4°C for six weeks also retained full protective activity at 12.5 and 2.5nM (Fig 1C, right).

Homeoprotein internalization is driven by the 3rd helix of the homeodomain (Derossi et al., 1994) and within this sequence mutating tryptophan (W) in position 48 of the homeodomain (HD) blocks internalization (Derossi et al., 1996). Accordingly, chEN2 internalization is abolished if the tryptophan and phenylalanine (W and F) residues at positions 48 and 49 of the HD are changed to serine and arginine (chEN2SR) residues, respectively (Joliot et al., 1998). Wild type mEN1, hEN1 and chEN2 provided 100% protection against H_2_O_2_ oxidative stress while no protection was observed by chEN2SR (Figure 1D), demonstrating that cell internalization is necessary for protection.

In addition to transcription, EN1 and EN2 also regulate protein translation (Brunet et al., 2005; Alvarez-Fischer et al., 2011; Stettler et al., 2012). Glutamate at position 50 of the homeodomain does not modify internalization but is necessary for high affinity DNA binding and transcriptional activity (Le Roux et al., 1993). To determine if ENGRAILED protective activity depended on transcription, hEN1 with a glutamine to alanine mutation at position 50 (hEN1Q50A) was produced. Figure 1D illustrates that, in contrast with wild-type hEN1, incubation with hEN1Q50A at the same concentrations provided no protection against oxidative stress. This demonstrates that EN1 protection against oxidative stress requires both internalization and high affinity DNA binding activity.

### Other HPs protect against oxidative stress

Mouse EN1, hEN1 and mouse or chEN2 are neuroprotective towards midbrain dopaminergic cells *in vitro* and *in vivo* (Sonnier et al., 2007; Alvarez-Fischer et al., 2011; Rekaik et al., 2015). Protection was also observed for OTX2 on DA midbrain cells *in vivo* and on RGCs *in vitro* and *in vivo* (Sonnier et al., 2007; Alvarez-Fischer et al., 2011; Torero et al., 2011; Rekaik et al., 2015). This raised the possibility that protective activity may be a property shared among a number of HPs. To verify if protection against oxidative stress is shared by several HPs from different classes, mOTX2, hLHX9, hGBX2 and chEN2 were compared to hEN1 in a single experiment with embryonic ventral midbrain neurons. Figure 2A demonstrates that the four HPs provided significant protection against 50mM H_2_O_2_ at 12.5nM. Only hLHX9 at 2.5nM failed to protect embryonic ventral midbrain neurons from oxidative stress-induced cell death.

**Figure 2.**
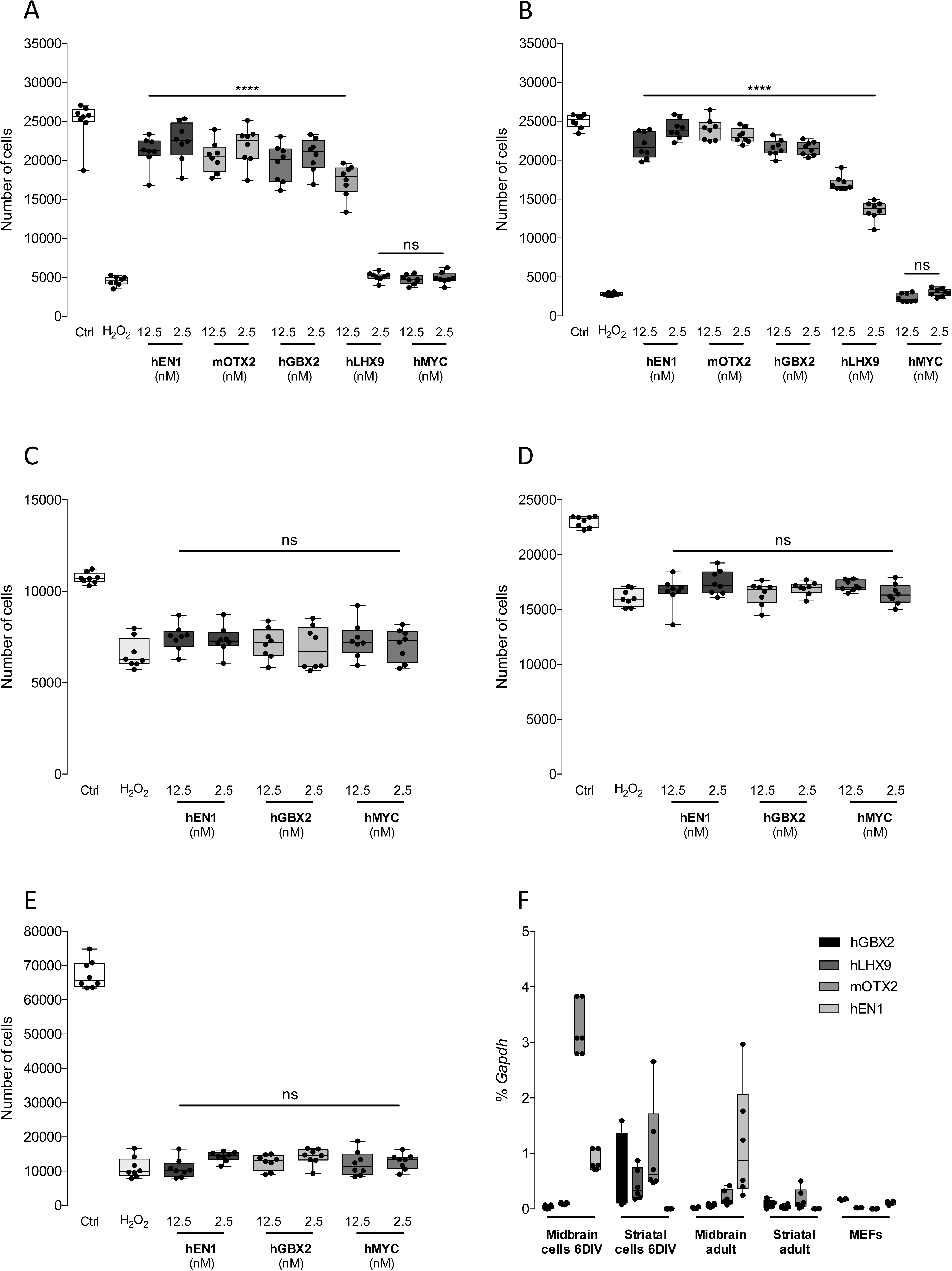
Homeoproteins protect embryonic neurons but not non-neuronal cells. A: hEN1, mOTX2, GBX2 and hLHX9 protect embryonic ventral midbrain cells against H_2_O_2_ oxidative stress while hMYC does not. B: hEN1, mOTX2, GBX2 and hLHX9 protect embryonic striatal neurons against H_2_O_2_ oxidative stress while hMYC does not. C: hEN1, hGBX2 and hMYC do not protect fibroblasts against H_2_O_2_ oxidative stress. D: hEN1, hGBX2 and hMYC do not protect HeLa cells against H_2_O_2_ oxidative stress. E: hEN1, hGBX2 and hMYC do not protect macrophages against H_2_O_2_ oxidative stress. F: RT-qPCR reveals expression of *GBX2, LHX9* and *OTX2* in embryonic striatum and *LHX9, OTX2* and *EN1* in ventral midbrain.

In contrast with the 4 HPs tested, a cell-permeable human MYC (hMYC) provided no protection (Figure 2A). In addition, protection by all HPs, but not hMYC, was also observed with striatal embryonic neurons (Figure 2B). This suggests that protection against oxidative stress may be specific to HPs with little neuronal subtype specificity. The fact that both striatal and midbrain neurons were protected and the absence of HP specificity led us to use hEN1 and hGBX2 to verify if they protected non-neuronal cells, including primary mouse fibroblasts (MEFs), HeLa cells and primary mouse macrophages. Figure 2 illustrates that although MEFs (Fig. 2C) and HeLa cells (Fig. 2D) are more resistant to oxidative stress than neurons (Fig. 2A,B) or macrophages (Fig. 2E) none of the non-neuronal cells are protected by the two tested HPs, or by cell-permeable c-MYC.

To verify if this large HP spectrum was related to unspecific HP expression in culture condition, we compared the expression of GBX2, LHX9, OTX2 and EN1 in 6 DIV cultures and adult tissues using RT-qPCR. Figure 2F illustrates that MEFs in culture express none of the HPs and that, in the embryonic cultures, GBX2 is expressed in striatal neurons only, LHX9 and OTX2 in midbrain and striatal neurons and EN1 in midbrain neurons only. For comparison, OTX2 is expressed in adult midbrain and striatum and EN1 in adult midbrain whereas LHX9 and GBX2 are barely expressed in the two structures (Fig 2F). These data demonstrate that a distinct HP can protect neurons that do not normally express it, two striking examples being the protection of striatal and midbrain neurons by EN1 and GBX2, respectively.

### Homeoproteins protect midbrain embryonic neurons against DNA damage

Oxidative stress causes a number of changes in cell physiology among which is the production of DNA breaks. In studies of neuroprotection of ventral midbrain neurons *in vivo*, Rekaik et al., (2015) observed that ENGRAILED reduces the number of anti-γH2AX-stained DNA damage foci induced in the nuclei of mDA cells exposed to 6-OHDA. To verify if this is also the case in the present *in vitro* conditions and for the 4 HPs studied, embryonic midbrain neurons were cultured for 6 DIV, treated with the mEN1 at a 2.5nM concentration for 24h and exposed for 1h to 100 μM H_2_O_2_. The cells were fixed and γ2HAX foci were revealed by immunocytochemistry in neurons identified by βIII tubulin labeling. Without H_2_O_2,_ neurons had only one or two γH2AX foci while H_2_O_2_ increased the number of foci about four-fold. Pretreatment with mEN1 reduced the number of γH2AX foci as illustrated in Figure 3A. As quantified in Figure 3B, the reduction in γH2AX foci was dose-dependent for mEN1, hLHX9, hGBX2 and mOTX2 at concentrations ranging from 2.3/3.3nM to 0.3/0.4Nm depending on the HP. Thus, each of the HPs tested protects neurons from oxidative stress, promotes their survival and reduces the level of DNA damage caused by H_2_O_2_.

**Figure 3.**
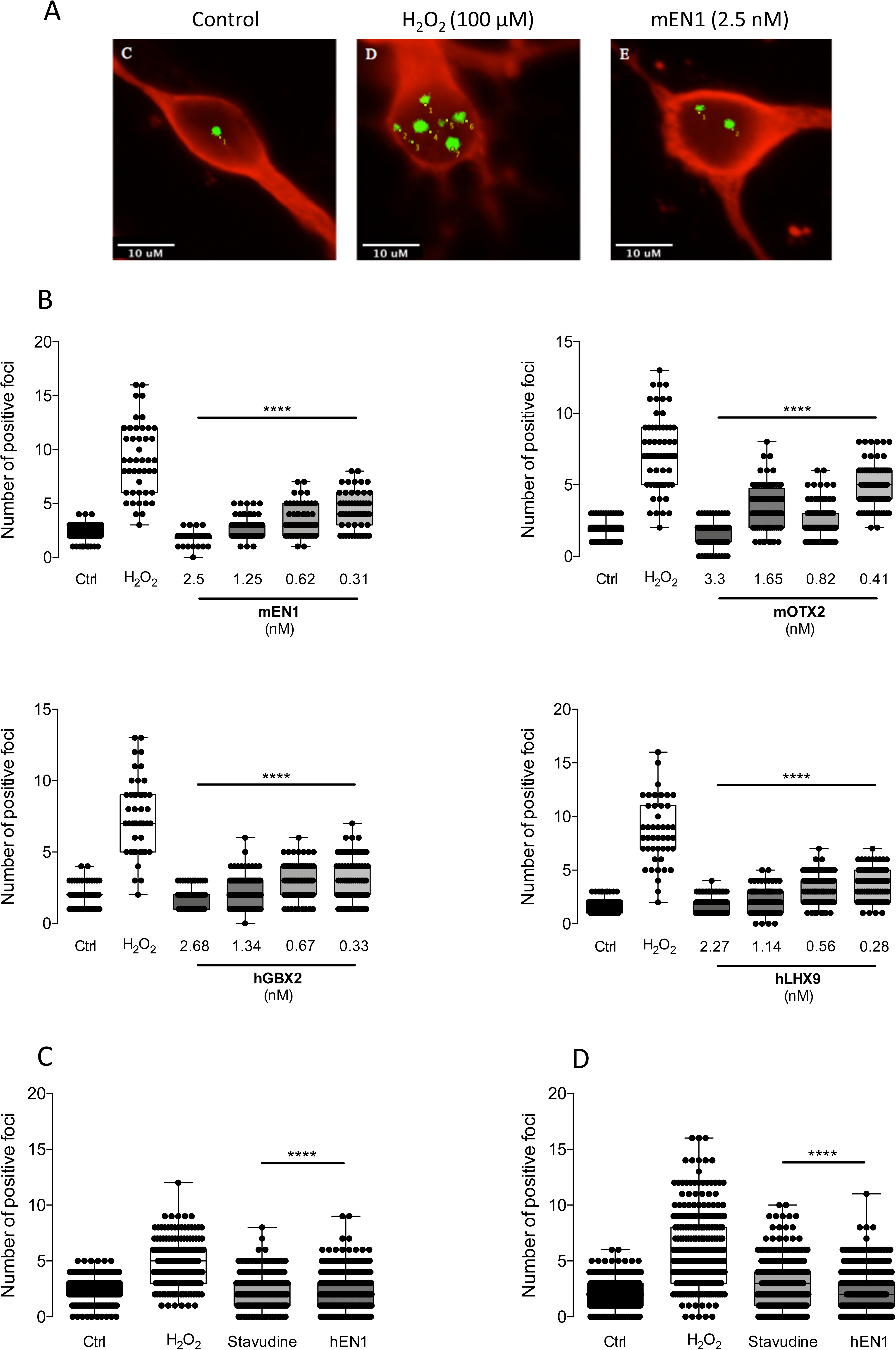
HPs reduce DNA breaks after H_2_O_2_. A: Cultures of E14.5 ventral midbrain neurons (red) untreated (Control) show few bright γH2AX foci (green) while those treated with 100μM H_2_O_2_ have numerous foci and those pretreated with mEN1 have only a few. B: Quantification of γH2AX foci. H_2_O_2_ increases the number of foci from about 1-2 per neuron to about eight. mEN1, mOTX2, hGBX2 and hLHX9 reduce the number of foci in a dose-dependent manner. C, D: Inhibition of reverse transcriptase activity protects against H_2_O_2_ oxidative stress in midbrain (C) and striatal neurons (D). In the control condition few γH2AX foci are observed in embryonic midbrain neurons while those challenged with 100μM H_2_O_2_ show multiple DNA damage foci. Pretreatment with 10µM stavudine or 2.5nM hEn1 completely blocks the formation of DNA damage foci.

Mobile element LINE-1 (L1) expression by midbrain neurons is increased by oxidative stress *in vitro* and *in vivo* and the endonuclease encoded by LINE-1 open reading frame 2 (ORFp2) is in part responsible for the breaks (Baudin de Thè et al., 2018). Accordingly, the protective activity of hEN1 is due to its ability to repress oxidative stress-induced LINE-1 overexpression (Baudin de Thè et al., 2018). Here, EN1 and the reverse transcriptase inhibitor stavudine used as a LINE-1 antagonist protected the oxidative stress–induced formation of DNA brakes in midbrain neurons. This led us to compare the effects of an overnight pretreatment by 12.5nM EN1 or 10μM and stavudine on the number of γH2AX foci in midbrain (Fig. 3C) and striatal cell cultures (Fig. 3D) following a one-hour incubation with 100µM H_2_O_2_. Figure 3C, D illustrates that 100μM H_2_O_2_ increased the number of γ-H2AX foci three-to four-fold in embryonic striatal and midbrain neurons compared to control (Figure 4). Stavudine at 10 µM significantly reduced the foci to the same extent as 12.5nM hEn1.

**Figure 4.**
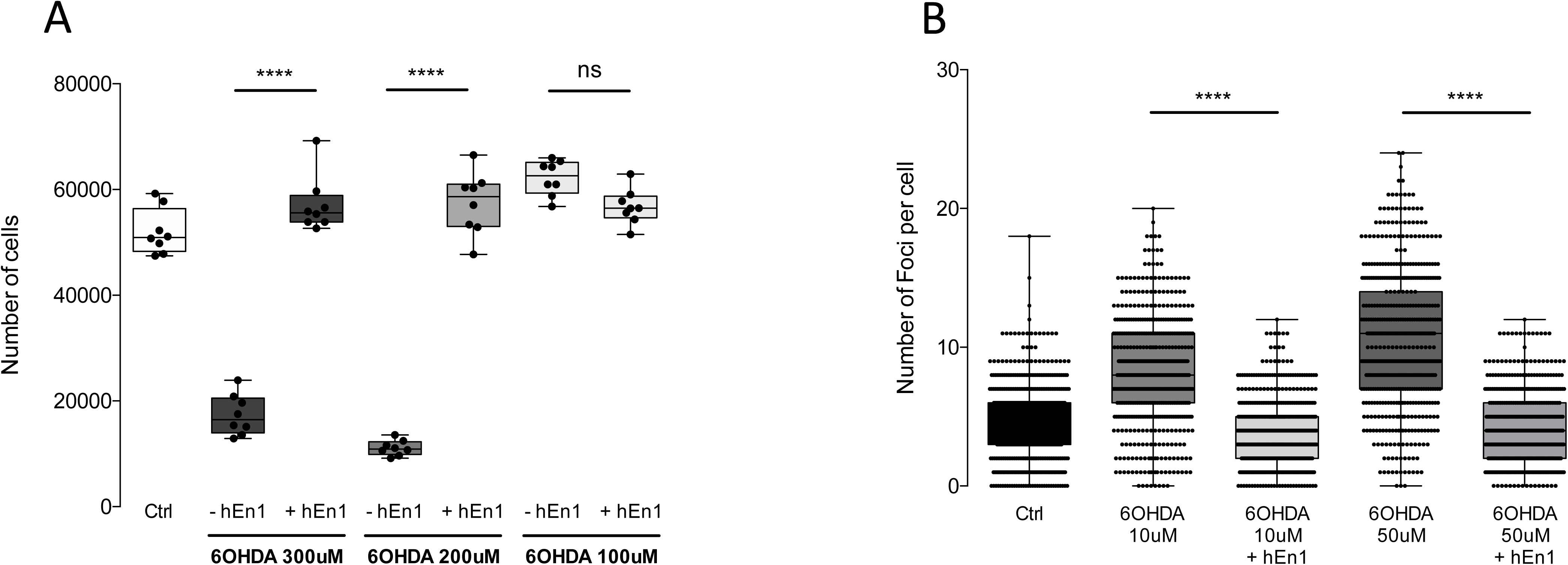
hEN1 protects immortalized human dopaminergic neuronal precursors, LUHMES cells against H_2_O_2_ oxidative stress. A: Three hundred and 200 μM 6-OHDA reduce the number of LUHMES cells surviving while 50nM hEN1 completely protects against this oxidative stress. B: Quantification of γH2AX foci. 6-OHDA increases the number of foci by 2-4 fold. Preincubation with hEN1 reduces the number of foci to the control level.

### hEN1 protects a dopaminergic cell line against 6-OHDA toxicity

ENGRAILED and OTX2 protect mesencephalic dopaminergic neurons *in vivo* against an oxidative stress induced by 6-OHDA (Rekaik et al., 2015). To verify if this is also the case in *vitro* and to follow the DNA-protection activity, we used immortalized human dopaminergic neuronal precursors, LUHMES cells, that express the DA transporter and are thus sensitive to 6-OHDA (as opposed to midbrain neurons in which the mDA neurons constitute a minority of the cell population). Figure 4 demonstrates that LUHMES cells are sensitive to the toxin at 200 and 300µM concentrations and entirely protected by a preincubation with 50nM hEN1. DNA breaks were also followed at lower 6-OHDA concentrations which induce breaks without provoking rapid cell death. As shown in Figure 4B, hEN1 reduces the number of breaks, confirming its protective activity in the 6-OHDA oxidative stress paradigm.

## DISCUSSION

Exogenous ENGRAILED protects DA neurons *in vitro* against MPP+ and rotenone and *in vivo* against 6-OHDA, MPTP, A30P α-synuclein and progressive degeneration associated with the loss of one EN1 allele (Sonnier et al., 2007; Alvarez-Fischer et al., 2011; Rekaik et al., 2015, Thomasson et al., 2019). OTX2 promotes the survival of adult dissociated RGCs *in vitro*, protects RGC *in vivo* against NMDA excitotoxicity and mDA neurons against 6-OHDA (Torero-Ibad et al., 2011; Rekaik et al., 2015). This similar pro-survival activity of two distinct transcription factors of different HP families led us to develop an *in vitro* assay to assess the ability of several HPs belonging to different classes to protect embryonic neurons against cell death and DNA damage caused by H_2_O_2_ oxidative stress. EN1, EN2 and GBX2 are members of the Antennapedia class, OTX2 belongs to the Paired class and LHX9 is part of the Lim class of HPs (Boncinelli, 1997).

The present results show that ENGRAILED internalization and high affinity DNA binding properties are necessary for its neuroprotective activity. This is in accord with previous results showing that when the WF at positions 85 and 86 in OTX2 (thus in positions 48 and 49 of its homeodomain) are mutated to YL, OTX2 loses its ability to be internalized and its neuroprotective activity for RGCs *in vitro* and *in vivo* (Torero-Ibad et al., 2011). The requirement for high affinity DNA binding suggests that survival activity implies transcriptional regulation and not by signal transduction of a cell surface receptor. This does not preclude activity at several other levels, including the regulation of protein synthesis or the maintenance of a healthy heterochromatin as demonstrated in studies on the protection of SNpc mDA neurons by ENGRAILED (Alvarez-Fischer et al., 2011; Stettler et al., 2012; Rekaik et al., 2015; Thomason et al. 2019). Whether these conclusions apply to all other HPs tested here is an open question.

DNA break-induced signaling such as the phosphorylation of the histone variant H2AX (γH2AX) is required for transcriptional elongation in healthy cells. In this case γH2AX accumulates at gene transcription start sites (TSSs) during Pol II pause release (Bunch et al., 2015). However, there are clear differences between the latter situation and γH2AX-marked DSBs induced by damaging conditions, including oxidative stress. In TSSs γH2AX accumulation is condensed within the transcribed units only and there is no spread outside the boundaries of the transcribed genes. In contrast, γH2AX accumulation due to DNA damage can spread over megabases in both directions from DSB sites. Here, the oxidative agents H_2_O_2_ and 6-OHDA significantly increased the number of γH2AX foci in embryonic neurons or LUHMES cells, respectively. Pretreatment with EN1, OTX2, GBX2 or LHX9 (embryonic neurons) or EN1 (LUHMES cells) prevented the formation of DSBs. Interestingly, another homeobox gene HOXB7 enhances non-homologous end joining DNA repair *in vitro* and *in vivo* (Rubin et al., 2007), providing additional support to the involvement of homeoproteins in DNA break repair.

Homeoproteins of different species (chicken, mouse and human) protect mouse embryonic neurons against oxidative stress, suggesting an evolutionary conservation of their protective activity that parallels their structure conservation (Banjeree-Basu and Baxevanis, 2001; Holland and Takahashi, 2005; Holland, 2013). The HPs tested here were all effective on neurons originating from the mesencephalon and telencephalon, two structures of different ontogenetic origins, thus expressing different repertoires of developmental genes. ENGRAILED and OTX2 expressed in the midbrain provide protection to striatal neurons and, conversely, GBX2 and LHX9 that are expressed in striatum are effective in providing protection to ventral midbrain neurons. These results raise the possibility that neuroprotective activity may be common to HPs in a non-region-specific manner. Interestingly, cell-permeant MYC, a transcription factor of the Basic Helix-Loop-Helix family (bHLH) with major roles in cell cycle progression, apoptosis and cellular transformation showed no neuroprotective effect against oxidative stress induced by H_2_O_2_.

In contrast with their neuroprotective activity for terminally differentiated non-proliferating embryonic neurons and the LUHMES immortalized human dopaminergic neuronal precursor cell line, none of the HPs tested was able to protect HeLa cells, primary macrophages or primary fibroblasts from H_2_O_2_ oxidative stress. Because all tests on non-neuronal cells were done in the presence of the Ara-C mitotic inhibitor and because the LUHMES cells are protected by hEN, it is unlikely that the absence of protection in non-neuronal cells is only due to their proliferative status. A possible explanation, based on the importance of hEN1 in mDA neurons in chromatin remodeling (Rekaik et al., 2015) is that the chromatin structure of the proliferative cells tested here is sensitive to HP expression. Alternatively, but not mutually exclusive, co-factors required of HP protection might not be available in these non-neuronal cells. Finally, oxidative stress increases LINE-1 expression and retrotransposition events increasing DNA damage. ENGRAILED reduces DAergic neurodegeneration by repressing LINE-1 expression *in vivo* (Blaudin de Thè et al., 2018). The results here extend the protective effects of the reverse transcriptase inhibitor, stavudine, to embryonic midbrain cells and striatal cells *in vitro*.

All in all, our results show that EN1, EN2, OTX2, GBX2 and LHX9 representing three different classes of HP transcription factors can protect embryonic cultured neurons from two ontogenetically diverse brain regions against H_2_O_2_-induced oxidative stress. The similar neuroprotection by ENGRAILED proteins from different species (i.e. chicken, mouse and human) demonstrates a strong evolutionary conservation of this activity. The ENGRAILED genes of vertebrates and insects arose as independent duplication of an ancestral EN gene (Dolecki and Humphreys, 1988) and this duplication occurred after the divergence of echinoderms and vertebrates but prior to the divergence leading to birds and mammals some 310 million years ago (Benton, 1993; Logan and Joyner, 1989; Kumar and Hedges, 1998). This suggests that ENGRAILED neuroprotective activity arose prior to the separation between birds and mammals. More strikingly, HPs can compensate between classes as shown by the fact that OTX2 of the PRD-class is able to compensate for EN1-dependent neuronal loss in vivo, even though ENGRAILED belongs to the ANTP-class (Di Giovannantonio et al., 2013). Thus, neuroprotective activity arose before the divergence of Antennapedia and paired classes of HPs. The early emergence of HP protection and the selective pressure to mantain it over hundreds of millions of years underscores the importance of this HP activity.

## Acknowledgements

We thank Marion Ruinart de Brimont, Yoko Arai, Bilal Mahzar, Raoul Torero-Ibad, Alain Joliot, Jessica Apulei and Ariel A. Di Nardo for producing some of the proteins used.

